# Standing genetic variation underlies divergence of nicotinic acetylcholine receptor subunits among cryptic species of the *Anopheles gambiae* complex

**DOI:** 10.64898/2025.12.07.692827

**Authors:** Caroline Fouet, Desiree E. Rios, Fred A. Ashu, Matthew J. Pinch, Cesar A. Hernandez, Marilene M. Ambadiang, Colince Kamdem

## Abstract

Arthropod species differ in insecticide susceptibility, yet how pre-existing polymorphism at target sites shapes variable responses within and among populations remains poorly understood. Recently diverged taxa provide ideal systems to test how target-site divergence modulates species sensitivity. Using whole-genome sequencing data from 573 mosquitoes representing six cryptic species of the *Anopheles gambiae* complex, we analyzed standing genetic variation across all 11 nicotinic acetylcholine receptor (nAChR) subunit genes to establish a baseline for natural diversity before the large-scale deployment of nAChR-targeting insecticides in Africa. We detected no previously reported resistance alleles from agricultural pests and found no evidence of selective sweeps or loss-of-function mutations across the nAChR gene family. Patterns of polymorphism were consistent with strong purifying selection. Most nonsynonymous variants were rare, predicted to be tolerated by SIFT (score ≥ 0.05), present almost exclusively in heterozygotes, and occurred outside ligand-binding and transmembrane domains. However, the α6 subunit exhibited relaxed constraint, with two high-frequency substitutions (I198M and D202E) that defined haplotypes segregating by species. The derived alleles represented ancient polymorphisms, showed evidence of introgression, and were fixed in populations with reduced larval susceptibility to spinosad. Our findings show that modest standing variation can shape divergence at insecticide target sites within a highly constrained gene family and underscore the need to monitor interspecific variation during the deployment of nAChR-targeting insecticides.

## 1. Introduction

Chemicals that interfere with neural signaling in insects form the basis of many insecticide classes used in agriculture and public health (Coats 2012). Resistance often arises from mutations that reduce the binding affinity of an insecticide to its molecular target, which lowers sensitivity in exposed populations (Georghiou 1972; McKenzie 1996; Hemingway and Ranson 2000). As a result, significant research has focused on detecting and functionally characterizing mutations within ligand binding domains or allosteric regions of insecticide target sites (ffrench-Constant et al. 1993; Martinez-Torres et al. 1998; Lenormand et al. 1999; Du et al. 2005; Platt et al. 2015; Assogba et al. 2016; Donnelly et al. 2016; Ranson and Lissenden 2016; Labbé et al. 2017; Grau-bové et al. 2020). More recently, the growing availability of genomic data has enabled the detection of positive selection at insecticide target sites, providing a common approach to identify potential resistance alleles (Jones et al. 2012; Faucon et al. 2015; Barnes et al. 2017; Kamdem et al. 2017; Miles et al. 2017; Fouet et al. 2018; Grau-bové et al. 2020; The Anopheles gambiae 1000 Genomes Consortium et al. 2020; Lucas et al. 2023; Boddé et al. 2025). However, selective sweep approaches (Maynard-Smith and Haigh 1974) provide only retrospective evidence of resistance evolution because the genomic signatures accumulate after selection, limiting their usefulness for real time surveillance. By contrast, dissecting standing genetic variation, in target site genes can provide essential early-warning information for resistance management. Such variants have been documented at major resistance loci, including *kdr* mutations in the voltage-gated sodium channel (Martinez-Torres et al. 1998) and *rdl* mutations conferring dieldrin resistance (Thompson et al. 1993), where alleles persisted in natural populations and facilitated rapid resistance evolution once insecticide pressure resumed (Curtis et al. 1998; Macoris et al. 2018; Grau-bové et al. 2020). Recently, new vector control insecticides with novel target sites have been approved, making it timely to characterize baseline variation before their widespread use.

Nicotinic acetylcholine receptors (nAChRs) are ligand gated ion channels that mediate rapid excitatory neurotransmission at postsynaptic membranes (Thany 2010). In arthropods, these receptors are the molecular targets of insecticide classes such as neonicotinoids, butenolides, and spinosyns, which act on overlapping but distinct binding sites (Jones and Sattelle 2010; Ihara et al. 2017; Matsuda et al. 2020; Perry et al. 2021; Komori et al. 2023). A functional receptor unit is composed of five subunits arranged around a central ion channel. Insects possess 10 to 12 nAChR subunit genes that encode α and non α subunits whose combination shape receptor pharmacology (Jones et al. 2005; Sattelle et al. 2005; Jones et al. 2006; Liu et al. 2013; Lu et al. 2022). Each subunit contains four transmembrane domains and six characteristic extracellular loops (A–F) that determine ligand specificity (Jones et al. 2005; Yao et al. 2008; Jones and Sattelle 2010; Komori et al. 2023). Although nAChRs are highly conserved in arthropods due to their essential roles in locomotion, circadian rhythm, learning, memory, courtship, sleep, longevity and immunity (Fayyazuddin et al. 2006; Somers, Luong, Mitchell, et al. 2017; Wang et al. 2024), point mutations within critical domains of α or β subunits have often been implicated in resistance to neonicotinoids and spinosyns in agricultural pests (Liu et al. 2005; Bass et al. 2011; Puinean et al. 2013; Chen et al. 2017; Yin et al. 2024; Cheng et al. 2025). Importantly, even in the absence of direct selection pressure, standing variation can shape receptor specificity and baseline susceptibility. For example, the limited sensitivity of ticks to neonicotinoids compared with other arthropods has been attributed to a conserved residue (Arg81) in the loop D binding region of the tick nicotinic acetylcholine receptor β1 subunit (Erdmanis et al. 2012). Neonicotinoids, butenolides, and spinosad are newly adopted classes of malaria vector control insecticides that target nAChRs, but their large-scale use in public health programs remains limited and they currently exert little to no selection pressure (WHO 2025). This provides a unique opportunity to explore genetic diversity in nAChRs among vector species and assess the role of standing variation in receptor divergence and ligand affinity.

The *Anopheles gambiae* complex provides an ideal model for studying the evolution of nAChRs in the absence of insecticide pressure. This cryptic species complex diverged relatively recently and now comprises taxa that occupy distinct ecological niches and display behavioral variability (Davidson and Jackson 1962; Coluzzi et al. 1977; Coluzzi et al. 1979; Coetzee et al. 2013; Barrón et al. 2019). Despite the absence of widespread exposure to nAChR-targeting compounds, interspecific differences in susceptibility to neonicotinoids have already been documented (Chouaibou et al. 2019; Dagg et al. 2019; Oxborough et al. 2019; Ashu et al. 2023; Ashu et al. 2024; Fouet, Ashu, et al. 2024; Kamya et al. 2024; Tchigossou et al. 2024; Oruni et al. 2025). For example, some *An. gambiae* larvae can survive and emerge at concentrations of neonicotinoids that are lethal to its sibling species *An. coluzzii* (Ambadiang et al. 2024). Although enhanced detoxification partly explains this difference (Fouet, Pinch, et al. 2024; Tchigossou et al. 2024), the finding also suggests that standing variation may be shaping the evolution of target sites within the species complex. Comparative analyses of nAChR evolution among cryptic species of *An. gambiae* sensu lato can thus offer critical insights into the interplay between standing variation, interspecific divergence, and insecticide susceptibility.

Here we combined whole-genome data from the *Anopheles gambiae* 1000 Genomes Project with targeted resequencing of the α6 subunit and larval bioassays to investigate interspecific divergence in nAChRs in *An. gambiae* sensu lato (Neafsey et al. 2015; The Anopheles gambiae 1000 Genomes Consortium et al. 2020). Although nAChRs are strongly constrained by purifying selection, substitutions in α6 (I198M and D202E) segregate by species across the *Anopheles gambiae* complex. The derived alleles are fixed in *An. gambiae*, whose larvae are less susceptible to spinosad than those of *An. coluzzii*, which retains the ancestral alleles. These findings provide a baseline for monitoring nAChR variation as new insecticides are deployed and highlight how standing variation in neural receptor genes can contribute to interspecific divergence in target-site responses.

## 2. Materials and Methods

### 2.1 Whole-genome dataset

We analyzed genetic variation using the *Anopheles gambiae* 1000 Genomes Phase 2-AR1 dataset, which includes whole-genome sequences from 1,142 wild-caught mosquitoes sampled from 33 sites across 13 sub-Saharan African countries (Miles et al. 2017; The Anopheles gambiae 1000 Genomes Consortium et al. 2020). This dataset has been described in detail by The Anopheles gambiae 1000 Genomes Consortium et al. (2020). Briefly, sequencing was performed using 100-bp Illumina reads at an average depth of approximately 30X per individual. Reads were aligned to the *AgamP4* reference genome using BWA v0.6.2 (Li and Durbin 2009), and variants were called with the GATK UnifiedGenotyper v2.7.4 (McKenna et al. 2010). For downstream analyses, we retained a subset of 532 individuals identified as *An. gambiae* (n = 343) or *An. coluzzii* (n = 189), sampled from 10 countries representing the geographic diversity of sub-Saharan Africa (Supplementary Table S1).

The *An. gambiae AgamP4* genome assembly contains 11 genes encoding nAChR subunits (9 α and 2 β) (Fig. 1A) (Jones et al. 2005). Exon coordinates for each gene were retrieved from the *AgamP4* genome annotation (GFF) using accession numbers (Giraldo-Calderon et al. 2015). Using bcftools v1.21 (Li et al. 2009), we extracted variants from the VCF files of the 532 selected individuals, retaining only those within exon regions and with a minor allele frequency (MAF) > 0.01 across each nAChR gene. We also analyzed whole-genome sequencing data from four species in the *Anopheles gambiae* complex—*An. arabiensis* (n = 12), *An. quadriannulatus* (n = 10), *An. melas* (n = 9), and *An. merus* (n = 10)—and extracted genotypes for α6 I198M and D202E from the corresponding VCFs (Neafsey et al. 2015).

**Figure 1.**
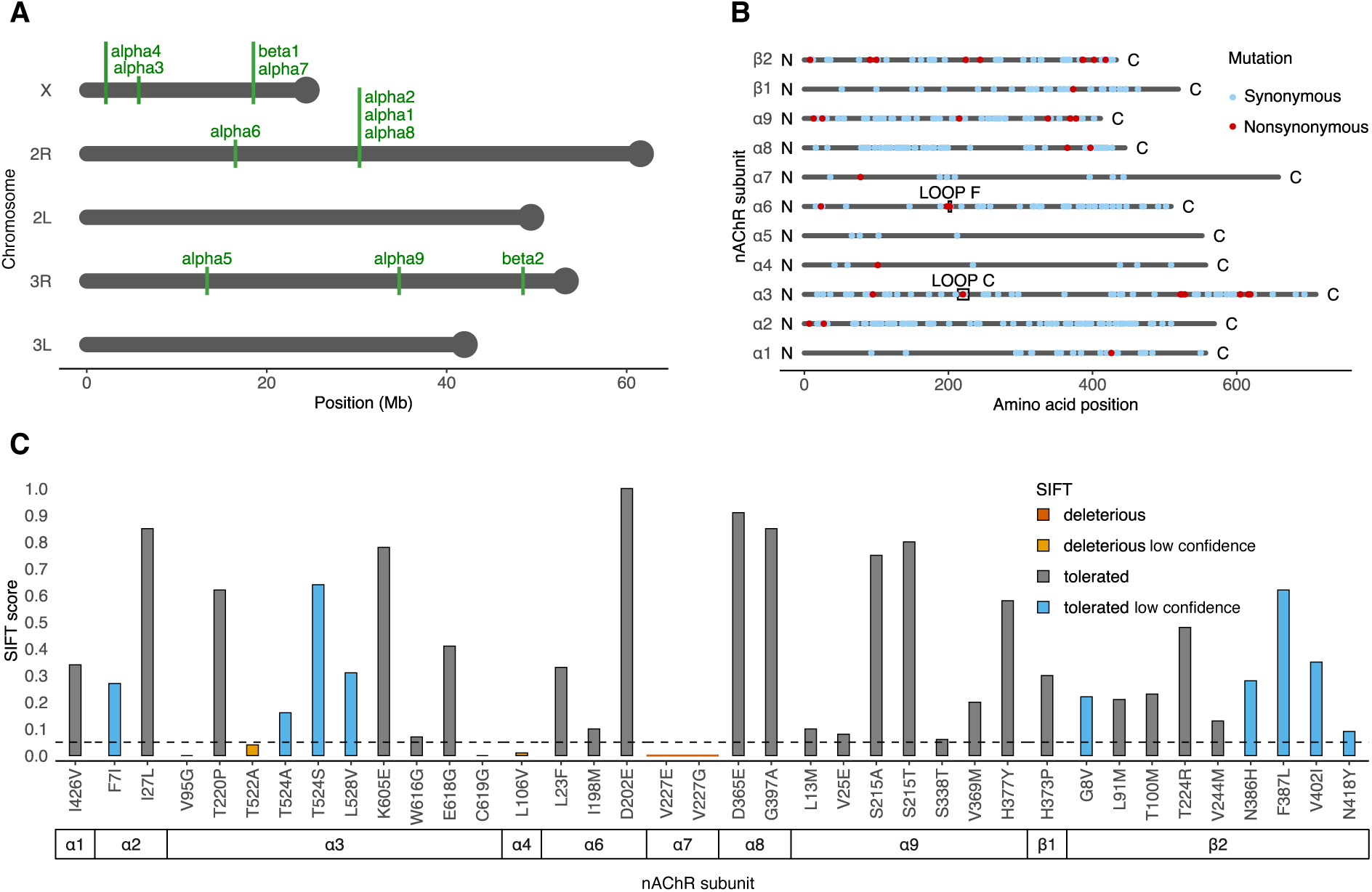
**Genomic locations, mutational landscape, and SIFT predictions for nAChRs in *An. gambiae* and *An. Coluzzii*** (A) Physical map of the 11 nAChR subunit genes along the five chromosome arms in the *Anopheles gambiae* PEST reference (*AgamP4*). (B) Positions of all detected mutations per subunit. Nonsynonymous variants that fall within ligand-binding loops (A–F) are highlighted. (C) SIFT scores for all nonsynonymous variants across nAChR subunits. Variants with SIFT < 0.05 (dashed horizontal line) are classified as predicted deleterious and ≥ 0.05 as tolerated. Predictions flagged as low confidence are indicated.

### 2.2 Variant annotation

We annotated exonic variants with SnpEff v4.3 (Cingolani, Platts, et al. 2012) using the *An. gambiae* reference database to classify variants by predicted functional impact across each nAChR subunit. Predicted effects on protein coding sequences were classified into three categories of impact as defined in SnpEff: (i) low impact: changes expected to have little or no influence on protein activity, here referred to as synonymous (e.g., synonymous variant, start retained variant, stop retained variant, 5ʹUTR variant, and 3ʹUTR variant); (ii) moderate impact: alterations that may affect protein function without completely disrupting it, here referred to as nonsynonymous (e.g., missense variant, inframe insertion, inframe deletion, and splice region variant) and (iii) high impact: mutations predicted to cause major disruption of the protein, here referred to as loss-of-function (e.g., stop gained, frameshift variant, splice acceptor variant, splice donor variant, and start lost). Annotation information was extracted from the annotated VCF files using SnpSift (Cingolani, Patel, et al. 2012). We screened the annotated data to identify nonsynonymous variants occurring within the predicted coordinates of the ligand binding loops (A–F) and the four transmembrane domains (TM1–TM4) (Jones et al. 2005). Finally, in addition to SnpEff impact categories, we annotated nonsynonymous variants in nAChR subunits with the Ensembl Variant Effect Predictor (VEP, v115) (McLaren et al. 2016), extracting SIFT (Sorting Intolerant from Tolerant) predictions and scores. Variants with SIFT < 0.05 were classified as deleterious and all others as tolerated (McLaren et al. 2016). Since VEP reports predictions per transcript, only the minimum SIFT score (i.e., the most deleterious prediction) was retained for each nonsynonymous variant among isoforms. Predictions flagged by VEP as low confidence were labeled accordingly.

### 2.3 Genetic diversity and divergence in nAChRs

To assess genetic diversity across nAChRs, we calculated the observed heterozygosity as the frequency of heterozygous genotypes for each subunit separately in *An. gambiae* and *An. coluzzii*. We compared heterozygosity at the gene level between synonymous and nonsynonymous mutations within each species. We also determined the proportion of exclusively heterozygous mutations, that is, the percentage of sites exhibiting heterozygous genotypes but no homozygous mutant individual.

To examine genetic differentiation, we extracted synonymous and nonsynonymous SNPs with MAF > 0.01 from the merged, annotated VCF of 532 individuals for each nAChR gene. We then converted these data to *genind* objects using the adegenet package (v2.1.10) (Jombart 2008). Nei’s genetic distances (Nei 1972) were subsequently calculated, and population structure was visualized using neighbor-joining trees (Saitou and Nei 1987). To quantify divergence between *An. gambiae* and *An. coluzzii*, we calculated relative genetic differentiation (*F*_ST_) per gene using Hudson’s estimator in PLINK v2.0 (Hudson et al. 1992; Chang et al. 2015). *F*_ST_ values were computed for all synonymous and nonsynonymous sites separately, and median *F*_ST_ values per gene were used to assess overall differentiation between *An. gambiae* and *An. coluzzii*. Finally, for each nonsynonymous mutation, we tested whether the mutant-allele frequency differed between species using Fisher’s exact test on allele counts.

### 2.4 Scanning for signatures of positive selection near α6 D202E

To test for signatures of a selective sweep around α6 D202E, we centered a ±200 kb window on the SNP (400 kb total) and computed extended haplotype homozygosity (EHH) decay from the mutation for the wild-type (D) and mutant (E) alleles on phased haplotypes (Sabeti et al. 2002). We pooled samples across species and grouped mutant (E/E, n = 322) and wild-type (D/D, n = 189) individuals. From the merged VCF, we extracted all biallelic SNPs within the ±200 kb window around D202E and phased genotypes with PHASE v2.1 (Stephens et al. 2001). We computed extended haplotype homozygosity (EHH) as a function of distance from the D202E codon for each allele using *ehh_decay* in scikit-allel (Miles et al. 2024). We compared the EHH decay curves between the mutant and wild-type groups to test for signals of positive selection linked to each haplotype.

### 2.5 Topology of α6 I198M and D202E

We localized the high-frequency substitutions I198M and D202E and quantified their proximity to the putative spinosad allosteric site. The receptor model was the AlphaFold-predicted monomer of *Anopheles gambiae* α6, with residue indices shifted by +19 to match coding-sequence numbering (AlphaFold 179 → 198; AF183 → 202). To approximate the spinosyn-binding site at the TM2–TM3 junction, we defined a centroid reference point as the weighted center of mass of three structural elements: the TM2–TM3 loop (0.6), the first three residues of TM3 (0.3), and the last two residues of TM2 (0.1). TM3 was identified by sequence match to the conserved motif YFNCIMFMVASSVVLTVVVLNY (Jones et al. 2005). TM2 was defined as the longest continuous α-helix beginning approximately 8–36 residues N-terminal to the start of TM3, and the TM2–TM3 loop was taken as the eight residues immediately preceding TM3.

Distances were measured in Å as (i) the minimal heavy-atom distance from each variant residue to TM3 and (ii) the Cα-to-centroid distance for residues I198 and D202. All selections, centroid calculations, and measurements were performed in PyMOL v2.6 (Schrödinger and DeLano 2020).

### 2.6 Distribution of α6 haplotypes in *Anopheles*

To examine the cross-species distribution of I198M and D202E, we retrieved α6 exon-7 protein sequences for 18 *Anopheles* species from VectorBase (including *An. arabiensis*, *An. melas*, *An. quadriannulatus*, and *An. merus*). The *Aedes aegypti* ortholog was used as an outgroup (Giraldo-Calderon et al. 2015). Sequences were aligned with ClustalW (Thompson et al. 1994) and visualized in ESPript (Gouet et al. 2003) to highlight residues 198 and 202.

Within the *An. gambiae* complex, allele frequencies for I198M and D202E were extracted from the *Anopheles gambiae* 1000 Genomes VCFs for *An. gambiae* (n = 342) and *An. coluzzii* (n = 189) across Africa. Linkage disequilibrium between the two sites was quantified as Lewontin’s Dʹ (Lewontin and Kojima 1960) using PLINK v1.19 (Purcell et al. 2007). For four additional members of the complex, genotype calls were obtained from Neafsey et al. (2015): *An. merus*, *An. melas*, *An. quadriannulatus*, and *An. arabiensis*. For each of the four species, we matched the VCF to its reference assembly using bcftools. We located α6 by mapping the *An. gambiae* α6 protein to the target genome with Exonerate (*–-model protein2genome*) (Slater and Birney 2005), which produced a GFF file. From this GFF file, we extracted a strand-aware α6 CDS and rebuilt the coding sequence by concatenating CDS blocks using samtools (Li et al. 2009) and custom Python scripts. The resulting α6 CDS FASTA for each species was aligned to the reference *An. gambiae* α6 sequence with MAFFT v7.526 (Katoh and Standley 2013), after which we queried the 3-bp codon windows at positions 198 and 202 using samtools/bcftools to obtain per-sample genotypes.

Finally, to test if local adaptation affects the distribution of I198M and D202E, we sequenced a section of α6 in populations from Cameroon. We previously designed a primer pair to amplify a region encompassing exons 6, 7, and 8 of the *An. gambiae* α6 subunit (Fouet, Pinch, et al. 2024). We used this primer pair to sequence *An. gambiae* and *An. coluzzii* populations sampled from diverse ecogeographic regions across Cameroon. This allowed us to study the frequency distribution of I198M and D202E in adult females collected from seven sites representing urban centers, the forest–savannah transition zone, coastal regions, and major agricultural areas (Fig. 4B). In parallel, we sequenced the same region in individuals from the *An. gambiae* Kisumu and *An. coluzzii* Ngousso laboratory colonies to assess whether I198M and D202E frequencies differ between wild populations and colonized strains reared under standard laboratory conditions. Genomic DNA was extracted from individual mosquitoes, and the target region was amplified using the designed primers. Amplicons were sequenced in both forward and reverse directions. Chromatograms were processed using Tracy v 0.7.8 (Rausch et al. 2020) to generate consensus sequences, which were then aligned to the *AgamP4* reference using MAFFT to identify the I198M and D202E mutations and calculate genotype frequencies. Species were identified as *An. gambiae* or *An. coluzzii* using molecular diagnostic PCR (Fanello et al. 2002).

### 2.7 Larval susceptibility to spinosad

We conducted standard larval bioassays (WHO 2005) to evaluate the sensitivity of nine *An. gambiae* and *An. coluzzii* populations to spinosad. Third-instar larvae were collected from seven sites in south of Cameroon and tested alongside the two laboratory colonies *An. gambiae* Kisumu and *An. coluzzii* Ngousso. Species identification of *An. gambiae* and *An. coluzzii* was performed using molecular diagnostic PCR on genomic DNA extracted from 100 larvae per population. Only populations unambiguously identified as *An. gambiae* or *An. coluzzii* were included in the larval bioassays. Spinosad is a naturally derived insecticide composed of two macrolide compounds: spinosyn A (∼85%) and spinosyn D (∼15%). It functions as an allosteric modulator of the nAChR α6 subunit and is approved as a mosquito larvicide (WHO 2025), although it has not yet been implemented in malaria control programs. We tested larval sensitivity to technical-grade spinosad (≥98% purity, Sigma-Aldrich) using ethanol-based stock solutions. Bioassays were conducted at two concentrations: 0.5 ppm, the WHO-recommended dose for vector control in drinking-water sources and containers (Hertlein et al. 2010; WHO 2010), and a lower dose (0.1 ppm).

Larvae were exposed to insecticides by transferring third-instar larvae in batches of 25 into 500-mL trays containing 200 mL of spinosad-treated water. Larvae reared in untreated laboratory water were used as control. Trays were covered with nets, and 10 mg of Tetramin fish food was added to each tray daily. The number of live third-and fourth-instar larvae and pupae was counted daily for six days. Dead larvae were equally counted and removed each day. Survival, mortality, and pupation rates were compared across populations and spinosad concentrations.

## 3. Results

### 3.1 The nAChR gene family is strongly constrained by purifying selection

Using whole-genome sequencing data from the *Anopheles gambiae* 1000 Genomes Project (The Anopheles gambiae 1000 Genomes Consortium et al. 2020), we examined genetic variation in nAChR genes in *An. gambiae* (n = 343) and *An. coluzzii* (n = 189) from 10 African countries. Across the 11 nAChR subunit genes in *Anopheles* (Jones et al. 2005), SnpEff annotations identified 36 nonsynonymous and 418 synonymous substitutions, and no predicted loss-of-function mutations (Cingolani, Platts, et al. 2012) (Table 1; Supplementary Material, Table S1). The gene family displayed signatures of purifying selection, consistent with the essential role of nAChRs in neural signaling (Fayyazuddin et al. 2006; Somers, Luong, Mitchell, et al. 2017; Wang et al. 2024).

**Table 1.** Distribution of synonymous and nonsynonymous variants across nAChR subunits in An. gambiae *and* An. coluzzii.

| Subunit | Protein length (AA) | CDS length (bp) | Synonymous mutations | Nonsynonymous mutations | Nonsynonymous mutation rate (%) |
| --- | --- | --- | --- | --- | --- |
| $\alpha 1$ | 557 | 1674 | 58 | 1 | 0.06 |
| $\alpha 2$ | 569 | 1710 | 89 | 2 | 0.12 |
| $\alpha 3$ | 710 | 2133 | 44 | 10 | 0.47 |
| $\alpha 4$ | 557 | 1674 | 6 | 1 | 0.06 |
| $\alpha 5$ | 552 | 1659 | 41 | 0 | 0.00 |
| $\alpha 6$ | 509 | 1530 | 33 | 3 | 0.20 |
| $\alpha 7$ | 658 | 1977 | 7 | 1 | 0.05 |
| $\alpha 8$ | 445 | 1338 | 19 | 2 | 0.15 |
| $\alpha 9$ | 411 | 1236 | 46 | 6 | 0.49 |
| $\beta 1$ | 519 | 1560 | 25 | 1 | 0.06 |
| $\beta 2$ | 433 | 1302 | 50 | 9 | 0.69 |

First, the subunit-encoding genes showed intolerance to loss-of-function mutations and a paucity of nonsynonymous substitutions in continental populations of *An. gambiae* and *An. coluzzii*. The distribution of the few missense mutations that occurred across subunits was uneven (Fig. 1A-B). Some subunits carried no nonsynonymous variants (α5) or only a single missense mutation (α1, α4, α7, and β1), whereas β2 and α3 harbored 9 and 10, respectively. Intermediate levels of variation were observed in α9 (6 nonsynonymous variants) and α2, α6 (3 nonsynonymous variants). Subunits with few mutations tended to harbor substitutions in the extracellular N-terminal region (α2, α4, α7) or intracellular C-terminal region (α8). By contrast, more variable subunits such as α9 and β2 displayed substitutions distributed along the protein sequence, though α9 showed clustering in both termini (Fig. 1B). The overall mutation rate, computed as the number of nonsynonymous substitutions per coding exon length, was very low, ranging from 0 in α5 to 0.0069 in β2 (Table 1).

Purifying selection was also reflected in the high conservation of critical functional regions including ligand-binding loops and transmembrane domains. To test whether missense mutations mapped to these regions, we examined nucleotide sequences corresponding to the four predicted transmembrane domains (TM1–TM4) and the six ligand-binding loops (A–F) in *An. gambiae* nAChRs (Jones et al. 2005). No missense mutations were detected within transmembrane domains, and only two of the 36 nonsynonymous substitutions occurred within ligand-binding loops (D202E in α6 loop F and T220P in α3 loop C) (Fig. 1B). This pattern indicates evolutionary constraint preventing the occurrence of amino acid substitutions across these critical regions. Moreover, none of the mutations previously reported to confer resistance in agricultural pests which typically involve modifications of ligand-binding affinity were observed in our dataset (Liu et al. 2005; Bass et al. 2011; Puinean et al. 2013; Chen et al. 2017; Yin et al. 2024; Cheng et al. 2025).

In addition to the absence of loss-of-function mutations, most amino-acid substitutions were predicted to have minimal functional impact. We annotated all nonsynonymous variants with the ensembl Variant Effect Predictor (VEP) (McLaren et al. 2016) and classified them as deleterious or tolerated using SIFT scores (Ng and Henikoff 2001). SIFT predicts whether an amino acid substitution is likely to affect protein function by analyzing sequence conservation from multiple alignments of homologous proteins using a score from 0 to 1. Variants with scores <0.05 are predicted deleterious and those ≥0.05 tolerated. Predictions derived from shallow or low-diversity alignments were flagged as “low confidence”. Across 38 missense variants identified by VEP in the *An. gambiae* nAChR family, 28 (74%) were predicted tolerated and 6 (16%) deleterious (Fig. 1C). Deleterious substitutions occurred in α3, α4, and α7. In α3, three positions scored as deleterious: V95G (0.00), T522A (0.04, low confidence), and C619G (0.00). In α7, two substitutions at the same residue were deleterious (V227G, 0.00; V227E, 0.00). α4 carried one deleterious variant (L106V, 0.01, low confidence). Outside these cases, missense variants were predicted tolerated on all subunits. α6 had three substitutions, all tolerated (L23F, 0.33; I198M, 0.10; D202E, 1.00), and the only nonsynonymous change in β1 (H373P, 0.30) was tolerated. The two high-frequency variants across all subunits (α6 I198M, 0.10; α6 D202E, 1.00) were tolerated, consistent with the persistence of substitutions with limited functional impacts under purifying selection.

Another important footprint of purifying selection was the overall reduction in genetic diversity, as expected for highly constrained gene families such as nAChRs (Fig. 2A). We quantified observed heterozygosity as the proportion of heterozygous genotypes at each synonymous and nonsynonymous site. Median heterozygosity across nAChR subunits was low (<10%) for both types of mutations in both species, with the lowest value (5.23%) recorded for nonsynonymous variants in *An. coluzzii*. Synonymous mutations showed a relatively even distribution of heterozygosity across subunits in both species (interquartile ranges (IQR): *An. gambiae* = 2.94 and *An. coluzzii* = 4.45), whereas nonsynonymous variants were more heterogeneous in *An. gambiae* (IQR = 8.94). In this species, five subunits (α4, α5, α6, α8, and β1) fell below 5% of nonsynonymous heterozygosity whereas α7 peaked above 35% (Fig. 2A). A two-sided paired Wilcoxon signed-rank test on heterozygosity revealed no significant difference in median values between species for either synonymous or nonsynonymous sites. The comparison between *An. gambiae* and *An. coluzzii* revealed *p* = 0.083 for synonymous variants and *p* = 0.308 for nonsynonymous variants. When comparing synonymous and nonsynonymous sites within species, *p*-values were 0.415 for *An. gambiae* and 0.476 for *An. coluzzii*. Despite the absence of significant differences in overall heterozygosity, the pronounced heterogeneity in nonsynonymous sites suggests stronger purging of deleterious alleles in specific subunits, in contrast to the more uniform and neutral evolution of synonymous variants.

**Figure 2.**
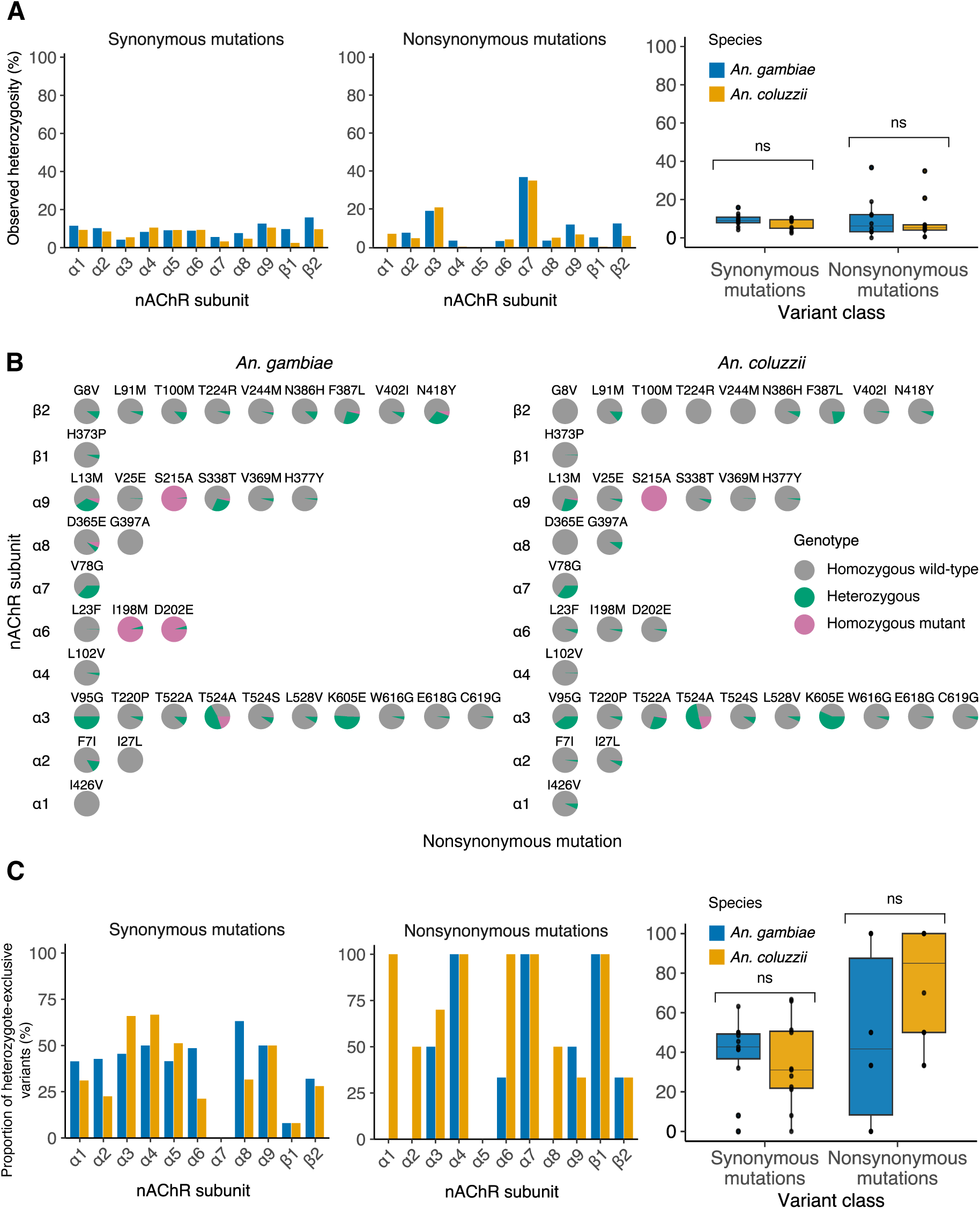
Heterozygosity and genotype frequencies for nAChR variants (A) Observed heterozygosity per subunit for synonymous and nonsynonymous mutations. Boxes show median and interquartile range. A two-sided paired Wilcoxon signed-rank test indicates no significant difference in median heterozygosity between species for either synonymous or nonsynonymous sites (ns, *p* > 0.05). (B) Genotype frequencies for all 36 nonsynonymous sites in *An. gambiae* and *An. coluzzii* populations across Africa. Mutations are grouped by subunit, and the frequencies of homozygous reference (wild-type), heterozygous and homozygous alternate (mutant) genotypes derive from whole-genome data (n = 532). (C) Fraction of heterozygote-exclusive variants (mutations observed exclusively in heterozygous genotypes) per subunit. Median and interquartile range are shown with a paired Wilcoxon test indicating no significant difference between species and variant class (ns, *p* > 0.05).

Finally, nAChR genes displayed a disproportionately high number of variants never observed in homozygous state supporting the presence of selective constraints and fitness costs associated with homozygosity at these loci. On average, the frequency of wildtype homozygotes was 78.08% in *An. gambiae* and 86.08% in *An. coluzzii* across nonsynonymous sites. Homozygous mutants were rare (3.62% on average in *An. coluzzii* and 9.55% in *An. gambiae*) for most variants (Fig. 2B). The rare exceptions among missense variants were S215A in α9 (∼100% homozygous mutants in *An. gambiae* and *An. coluzzii*) and the α6 substitutions I198M and D202E (95.63% for both in *An. gambiae*). Out of 36 SnpEff-annotated nonsynonymous mutations, 22 were exclusively found in heterozygotes and 7 showed heterozygotes outnumbering homozygous mutants. The proportion of exclusively heterozygous sites (i.e., variants with heterozygous genotypes but no homozygous individuals) was evenly distributed among synonymous variants at intermediate values consistent with genome-wide expectations (IQR: *An. gambiae* = 12.6% and *An. coluzzii* = 28.8%) (Fig. 2C). In contrast, nonsynonymous substitutions displayed heterogeneity across subunits, with several loci showing 100% exclusively heterozygous sites (IQR: *An. gambiae* = 79.2% and *An. coluzzii* = 50.0%). Within species, *An. gambiae* exhibited substantially greater dispersion for exclusively heterozygous mutations (12.6% for synonymous variants vs 79.2% for nonsynonymous sites, Δ (synonymous − nonsynonymous) = −66.6, 95% CI [−88.6, −44.7], *p* = 0.0071). In *An. coluzzii*, nonsynonymous mutations were enriched for exclusively heterozygous sites, with a median proportion 2.74 times higher than that of synonymous variants (paired Wilcoxon signed rank tests, *p* = 0.0144). The persistence of moderate-effect alleles in heterozygotes and their absence in homozygotes suggest that deleterious variants are maintained under mutation–selection balance (Haldane 1937) but are purged when homozygous, reinforcing the idea that nAChRs are subject to strong functional constraint.

### 3.2 Standing variation in α6 diverge between *An. gambiae* and *An. coluzzii*

We next asked whether, despite evidence of purifying selection, nonsynonymous variants segregate between *An. gambiae* and *An. coluzzii* populations at the continental scale and could contribute to species-specific divergence of nAChRs. To address this, we first constructed neighbor-joining trees for each subunit based on both synonymous and nonsynonymous variants. Levels of clustering varied across subunits. Subunits α3, α4, α5, and α9 showed minimal structuring by species or geography (Fig. 3A). β2 exhibited some differentiation, though not clearly aligned with species boundaries. In contrast, α1, α2, and α8 displayed distinct clustering into two groups corresponding to *An. gambiae* and *An. coluzzii*. The strongest segregation was observed for α6, α7, and β1, suggesting that these subunits may drive interspecific divergence in nAChRs.

**Figure 3.**
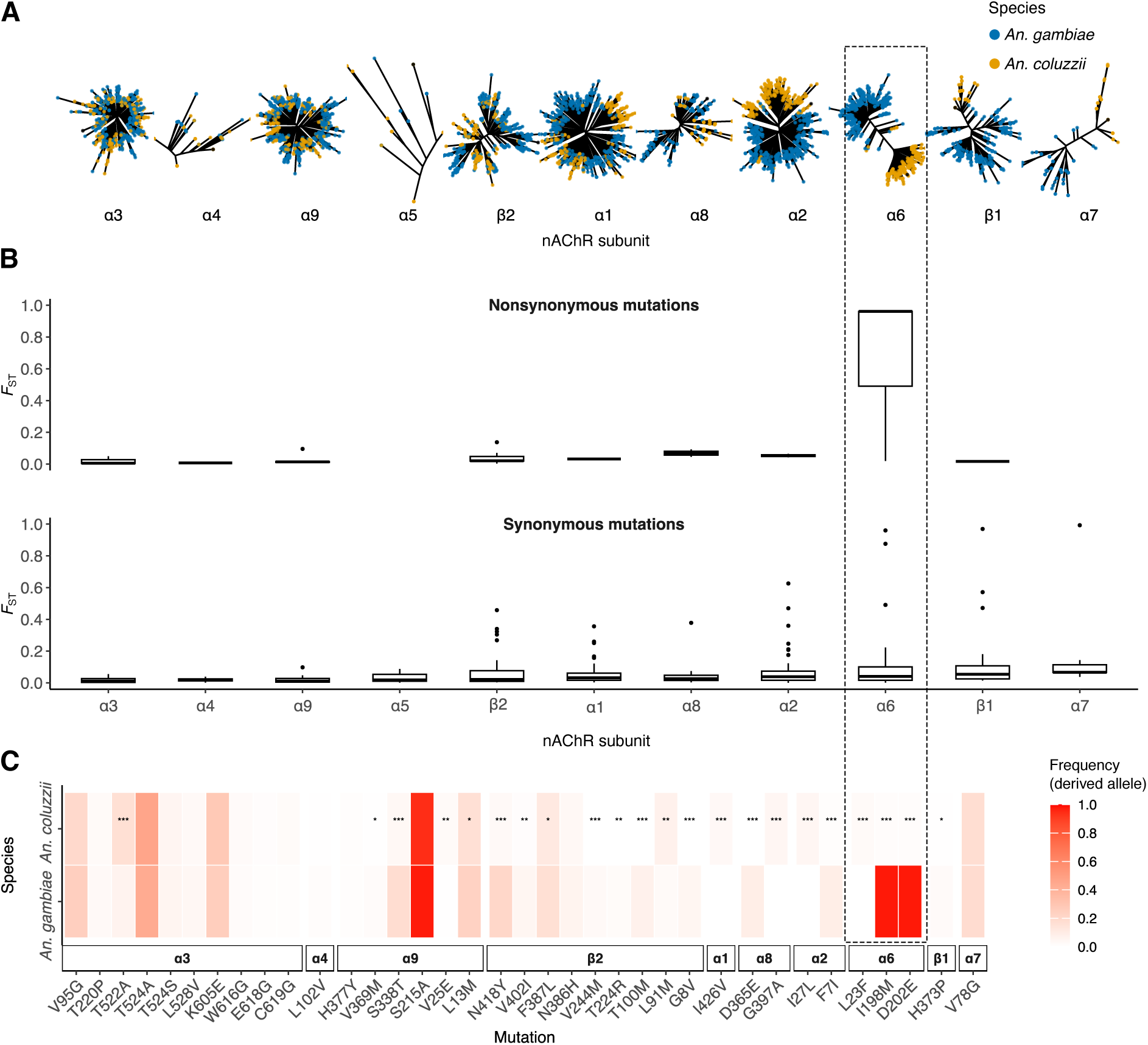
Divergence between *An. gambiae* and *An. coluzzii* across nAChR genes (A) Neighbor-joining trees per subunit built from genetic distances using nonsynonymous and synonymous mutations, highlighting α6, β1, and α7 as the most differentiated subunits. (B) Hudson’s *F*_ST_ per subunit for synonymous and nonsynonymous variants. Only α6 shows significantly high nonsynonymous differentiation between *An. gambiae* (n = 343) and *An. coluzzii* (n = 189) driven by the I198M and D202E substitutions. Nonsynonymous *F*_ST_ values are extremely low on other subunits consistent with shared polymorphism. (C) Heatmap of derived-allele frequencies for the 36 nonsynonymous SNPs in *An. gambiae* and *An. coluzzii*. Asterisks denote interspecific frequency differences by Fisher’s exact test: *p* < 0.05 (*), < 0.01 (**), < 0.001 (***).

**Figure 4.**
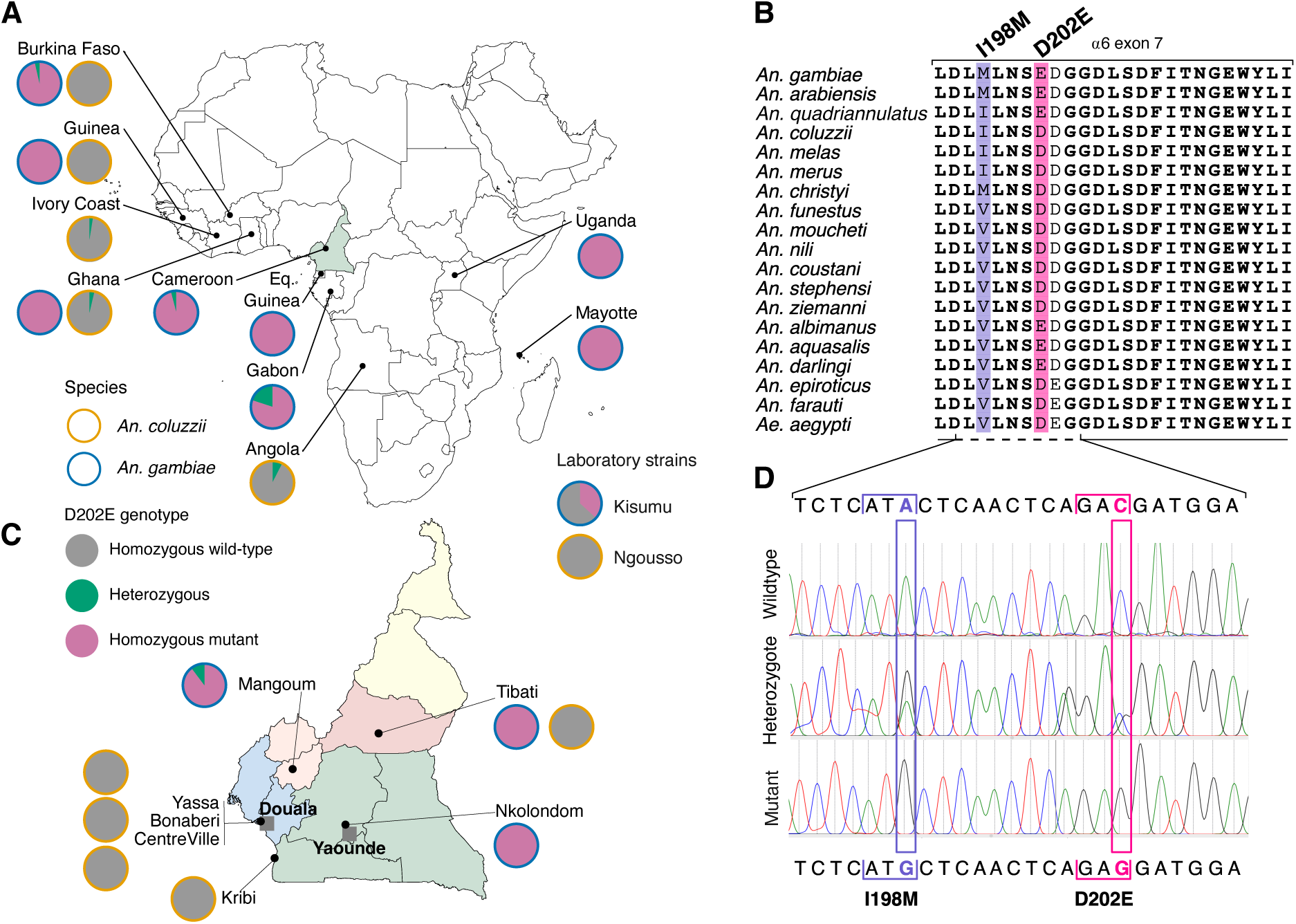
Geographic and phylogenetic distribution of α6 variants I198M and D202E (A) Country-level genotype frequencies for D202E in *An. gambiae* and *An. coluzzii* across Africa. D202E is in complete linkage with I198M (Lewontin’s Dʹ = 1), and the two distributions mirror each other. (B) Multiple alignment of α6 exon-7 protein sequences from 18 *Anopheles* species with *Aedes aegypti* as outgroup; residues 198 and 202 are highlighted. (C) Spatial distribution of D202E genotypes in Cameroon. The base map delineates major eco-geographic zones—equatorial/monsoon (green/blue/pink), tropical savanna (orange), and semi-arid (yellow). Sampling sites show species-specific segregation: *An. coluzzii* retains the ancestral D allele, whereas *An. gambiae* carries the derived E allele. (D) Representative Sanger chromatograms illustrating homozygous reference (I/D), heterozygous, and homozygous alternate (M/E) genotypes at positions 202 and 198.

We further quantified genetic differentiation by calculating *F*_ST_ separately for nonsynonymous and synonymous variants. Overall, *F_ST_* values for both types of mutation were low (median <0.05), consistent with the weak genome-wide divergence between the sister species (Fig. 3B) (della Torre et al. 2002; Miles et al. 2017). Nonetheless, several subunits including α6, α7, and β1 contained SNPs with *F*_ST_ values exceeding 0.8, and α2 harbored variants above 0.4. Closer examination revealed contrasting patterns between synonymous and nonsynonymous variants. Synonymous variants mirrored the tree-based clustering results, with α6, α7, and β1 emerging as the most differentiated subunits, characterized by higher median *F*_ST_ and an accumulation of outlier SNPs. In contrast, nonsynonymous variants identified α6 as the sole subunit exhibiting strong interspecific differentiation, suggesting a single region of functional divergence across the nAChR gene family between *An. gambiae* and *An. coluzzii*.

Despite their low frequencies, nonsynonymous mutations accumulated asymmetrically between species. *An. gambiae* contained more mutant alleles than *An. coluzzii*. Of the 36 missense variants, 22 showed significant frequency differences (Fisher’s exact test, *p* < 0.01), with 15 mutant alleles enriched in *An. gambiae* versus 7 in *An. coluzzii* (Fig. 3C). Especially in β2, one of the most variable subunits, all mutant alleles occurred at higher frequencies in *An. gambiae*. The two high-frequency nonsynonymous mutations in nAChRs (I198M and D202E in α6) were fixed in *An. gambiae* but absent from *An. coluzzii*.

Overall, while most nAChR subunits shared polymorphisms between species, significant divergence was concentrated in α6, α7, and β1. In α7 and β1, however, differentiation was driven by synonymous substitutions, as the single missense variant identified in each subunit was rare and showed no allele frequency difference between species. By contrast, two tolerated substitutions in α6 exon 7 (I198M and D202E) emerged as the principal drivers of functional divergence between *An. gambiae* and *An. coluzzii*. The rise in frequency of these amino acid changes supports the hypothesis that, contrary to *An. coluzzii*, *An. gambiae* α6 can tolerate non-deleterious mutations with limited fitness cost (Perry et al. 2007; Wang et al. 2016; Guillem-Amat et al. 2020; Perry et al. 2025).

### 3.3 α6 haplotypes recently emerged within the *An. gambiae* complex

Linkage disequilibrium analyses indicated that α6 D202E and I198M segregate together (Lewontin’s Dʹ = +1) and define two haplotypes in contemporary populations of *An. gambiae* and *An. coluzzii* at the continental scale. Across ten countries, the I198–202E haplotype was nearly fixed in *An. gambiae* but remained extremely rare in *An. coluzzii* (Fig. 4A). Among 343 *An. gambiae* samples, no homozygotes for the wild-type alleles (I198, D202) were detected, although 15 D202 heterozygotes (4.37%) were observed in Central and West Africa. Conversely, none of the 189 *An. coluzzii* samples carried the mutant alleles as homozygotes. Six heterozygotes (3.17%) were identified, all otherwise homozygous for the wild-type alleles. Consistent with the observed *F_ST_* values, these patterns indicate that the 198M–202E haplotype is restricted to *An. gambiae*, whereas I198–D202 is characteristic of *An. coluzzii*.

To test whether α6 haplotypes correlate with local adaptation at finer geographic scales, we examined their frequency distributions in *An. gambiae* and *An. coluzzii* across the main ecogeographic domains of Cameroon. Sequencing of exon 7 confirmed the partitioning of two haplotypes along species boundaries, consistent with continental-scale patterns (Fig. 4C-D). Among 161 field-collected *An. gambiae*, all individuals were homozygous for the derived alleles 198M and 202E, except four heterozygotes detected in Mangoum, an agricultural site. In contrast, all *An. coluzzii* individuals including all field populations as well the laboratory strain *An. coluzzii* Ngousso were homozygous for the wild-type alleles. In contrast, allele frequencies differed between field-collected and long-established laboratory colonies of *An. gambiae*. In the Kisumu strain, both mutant alleles occurred at substantially lower frequencies than in field populations (36.8% vs. 96.2% for 202E; Fisher’s exact test, *p* = 4.78 × 10⁻¹²). These findings suggest that while the continental and national distributions reflect interspecific differentiation rather than local adaptation, within-species variation persists in *An. gambiae*. The higher prevalence of the derived alleles in the wild is consistent with a selective advantage or relaxed purifying selection in natural settings, or with reduced fitness costs relative to long-term laboratory conditions.

The I198M and D202E substitutions, which are tightly linked in *An. gambiae* and *An. coluzzii*, segregate differently across other members of the *An. gambiae* complex. We identified α6 orthologs in four additional species, mapped positions 198 and 202 to their reference assemblies, and extracted genotypes from VCFs for 42 individuals (Neafsey et al., 2015). In *An. arabiensis*, the mutant allele M at 198 was fixed (12/12) and E at 202 was nearly fixed (11/12, one D/E). In *An. quadriannulatus*, mutation 198 remained wild-type (10/10 I), whereas the mutant allele E at 202 was nearly fixed (9/10, one D/E). In contrast, *An. melas* and *An. merus* carried only the wild-type alleles at both sites: I at 198 and D at 202 in all genotyped samples (Supplementary Table S2).

To visualize the distribution of I198M and D202E across *Anopheles* species, we aligned α6 exon-7 protein sequences from 18 taxa, using *Aedes aegypti* as outgroup (Fig. 4B). At α6-198, isoleucine (I) predominates within the *An. gambiae* complex while methionine (M) occurs on a shared haplotype in *An. gambiae* field populations and *An. arabiensis*. In more distant *Anopheles* outgroups, valine (V) is common at this site, consistent with isoleucine being ancestral within the complex. At α6-202, species of the complex are split in two groups: *An. gambiae*, *An. arabiensis*, and *An. quadriannulatus* carry glutamate (E), whereas *An. coluzzii*, *An. melas*, and *An. merus* retain aspartate (D). Most outgroups also carry D, supporting D as ancestral and E as derived. This patchy distribution, together with tight linkage between positions 198 and 202 in several species, is more consistent with introgression and/or retention of ancestral polymorphism than with multiple independent origins within the *Anopheles gambiae* complex.

### 3.4 No evidence of positive selection around D202E

To test whether the α6 locus has been subjected to selection pressures unrelated to insecticide use, we evaluated signals of positive selection associated with the I198M–D202E haplotype. Because the two mutations are in perfect linkage (D’ = 1), we used extended haplotype homozygosity (EHH) in a 400-kb window to explore signatures of selection centered on one of them (D202E). Across the 5-kb flanking regions on either side of the mutation, EHH decay was indistinguishable between mutant and wild-type chromosomes (Fig. 5A). In both populations, EHH values were > 0.95 at the D202E codon but dropped sharply to < 0.1 within 200 bp. Beyond this point, both traces remained close to zero, showing no evidence of extended haplotype structure as would be expected under a recent selective sweep (Maynard-Smith and Haigh 1974; Sabeti et al. 2002). This pattern is consistent with the absence of strong positive selection acting on either haplotype that differentiates *An. gambiae* and *An. coluzzii*.

**Figure 5.**
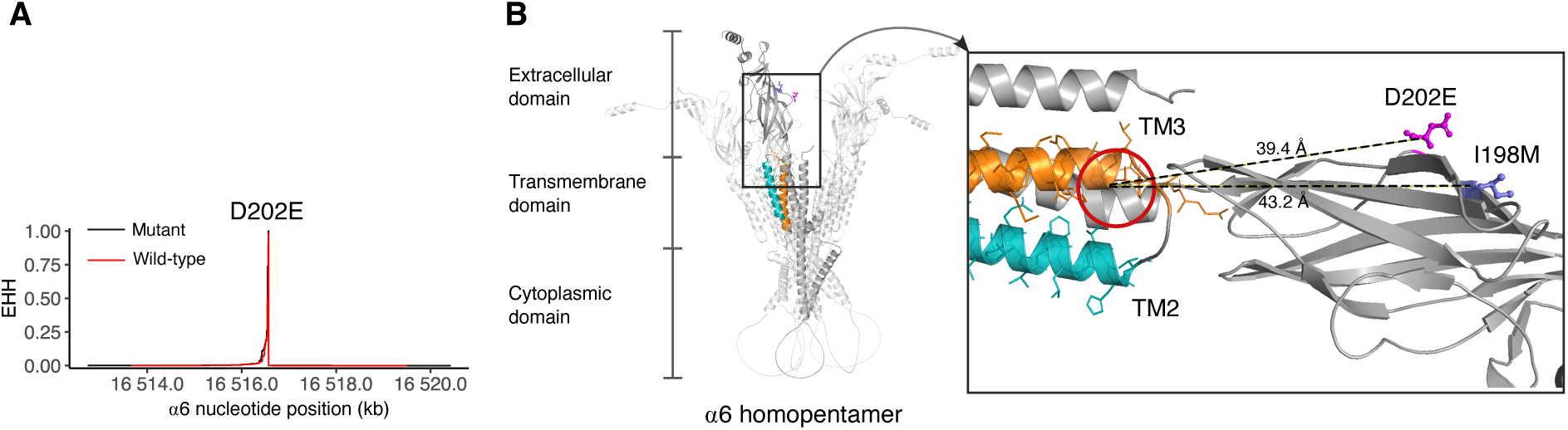
Haplotype homozygosity around α6 D202E and structural mapping of α6 substitutions (A) Extended haplotype homozygosity (EHH) decay from the α6 D202E core SNP for chromosomes carrying the ancestral D allele (wild-type) and the derived E allele (mutant) within a ±200 kb window. Genotypes were phased and EHH was computed as a function of distance from the core. Both allele classes show rapid EHH decay, indicating no evidence of a recent strong sweep centered on D202E. (B) Structural mapping of I198M and D202E in α6. Left: AlphaFold-based model of an α6 homopentamer with one subunit colored to highlight TM2 and TM3. Right: approximate spinosad allosteric site at the TM2–TM3 junction (orange), with Cα–centroid distances from I198 and D202 shown (∼39.4 Å and ∼43.2 Å, respectively).

### 3.5 Baseline susceptibility to spinosad differs between α6 haplotypes

In insects, resistance to spinosad is more frequently linked to alterations in the target site—the nAChR α6 subunit—rather than metabolic detoxification (Perry et al. 2007; Watson et al. 2010; Puinean et al. 2013; Perry et al. 2021; Perry et al. 2025; Sparks et al. 2025). Previous studies have shown that loss-of-function variants and missense mutations in the allosteric binding sites can confer resistance, and that α6 is a relatively plastic subunit, where mutations may carry low fitness costs (Perry et al. 2007; Baxter et al. 2010; Ureña et al. 2019; Guillem-Amat et al. 2020; Martelli et al. 2022; Perry et al. 2025). The *An. gambiae* Alphafold-predicted protein structure of α6 indicates that I198M and D202E mutations are located ∼40 Å from the putative spinosad allosteric binding site (Puinean et al. 2013) and are therefore unlikely to directly affect the binding affinity of spinosyns (Fig. 5B). Nevertheless, because insecticide sensitivity depends on complex interactions within the pentameric receptor that may not be captured by structural modelling of a single subunit, we directly tested the susceptibility of populations carrying different α6 haplotypes to spinosad. We assessed larval susceptibility in *An. gambiae* and *An. coluzzii* populations from Cameroon. Bioassays were conducted at two field-realistic doses (0.1 and 0.5 ppm) on third-instar larvae from six neonicotinoid-susceptible *An. coluzzii* populations including five filed populations and the Ngousso strain. In *An. gambiae*, the susceptible strain Kisumu was tested alongside a field population (Nkolondom) characterized by multi-resistance, including reduced susceptibility to neonicotinoids (Ashu et al. 2023; Ambadiang et al. 2024; Ashu et al. 2024; Fouet, Ashu, et al. 2024).

Survival analyses revealed interspecific differences. Across all six *An. coluzzii* populations, mortality was rapid and complete at both spinosad doses, with survival probabilities reaching zero by day 4–5 (Fig 6). In contrast, *An. gambiae* larvae from Nkolondom showed delayed mortality. At 0.1 ppm, a fraction of individuals survived until day 7 (25.9%), and at 0.5 ppm, survival probabilities were higher relative to *An. coluzzii* (log-rank test: Nkolondom vs Penja: χ² = 247.08, *p* < 2e-16; Nkolondom vs Combattant: χ² = 172.65, *p* < 2e-16). Pupation rates were consistent with mortality and survival. In *An. coluzzii*, pupation was absent at 0.5 ppm and limited to ∼30% at 0.1 ppm (Dibamba, 32% at day 2), whereas *An. gambiae* Nkolondom larvae continued to pupate throughout the assay, reaching cumulative pupation rates of 84% at 0.1 ppm and 12% at 0.5 ppm. In all control experiments without insecticide, 100% survival was obtained, and all larval populations reached the pupa stage within 72 h. These results indicate variability in baseline susceptibility to spinosad between species. In particular, the *An. gambiae* Nkolondom population—fixed for the derived 198M–202E haplotype—showed reduced sensitivity compared with *An. coluzzii* populations, which are fixed for the ancestral I198–D202 haplotype.

**Figure 6.**
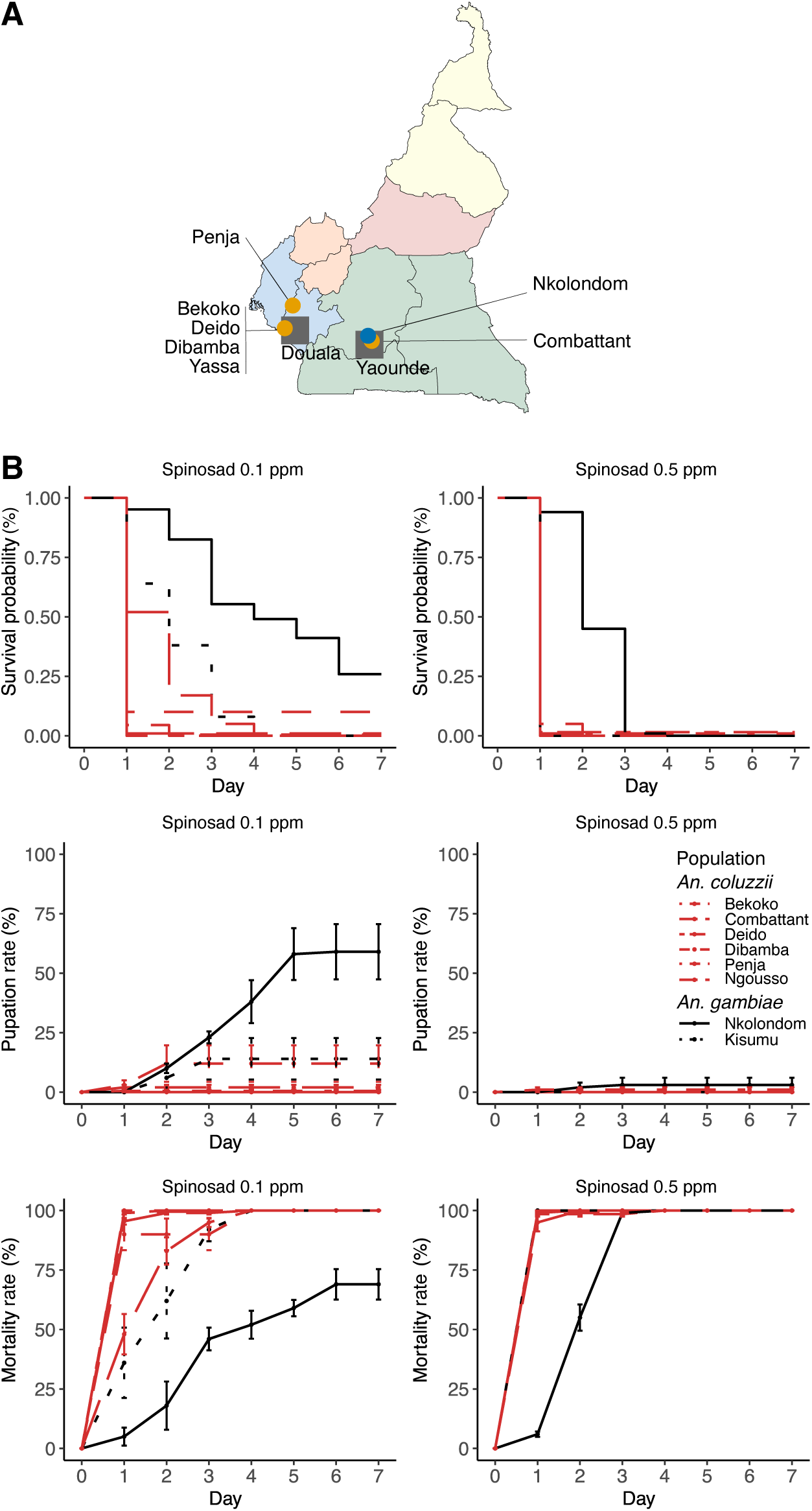
**Larval susceptibility to spinosad in *An. gambiae* and *An. Coluzzii*** (A) Sampling map showing larval collection sites in Cameroon: *An. coluzzii* (yellow) and *An. gambiae* (blue). (B) Seven-day assays on third-instar larvae exposed to spinosad (0.1 and 0.5 ppm). Panels report survival, pupation, and cumulative mortality for six field populations and two laboratory colonies (labels in panel). Survival differences between species at each concentration were evaluated with log-rank tests; *An. gambiae* exhibited higher survival (p < 0.05), consistent with higher pupation rates and lower mortality relative to *An. coluzzii*.

## 4. Discussion

### 4.1 Purifying selection with enhanced conservation of ligand-binding and transmembrane domains in nAChRs

Although studies have demonstrated that some insects can survive with impaired function of individual subunits such as α6 (Perry et al. 2007; Ureña et al. 2019; Guillem-Amat et al. 2020; Martelli et al. 2022; Mocchetti et al. 2025; Perry et al. 2025), disruptions to nAChR function generally impose substantial fitness costs, making these genes one of the most conserved among neuronal gene families (Somers, Luong, Batterham, et al. 2017; Ruel et al. 2019; Lu et al. 2022; Zhang et al. 2022; Zhang et al. 2022). Here we show that in *Anopheles* mosquitoes the integrity of nAChRs and their essential role in mediating fast excitatory neurotransmission are maintained by purifying selection as evidenced by the absence of loss-of-function mutations, the rarity of nonsynonymous variants and the conservation of ligand-binding and transmembrane domains. One of the anticipated consequences of purifying selection is that most strongly deleterious loss-of-function variants will be removed from the population or maintained at very low frequencies, thus preventing them from becoming common (Gorlov et al. 2006; MacArthur et al. 2012; Rausell et al. 2020). Our study highlights loss-of-function intolerance as the prime indicator of evolutionary constraints across nAChRs in *An. gambiae* and *An. coluzzii*. These genes are further constrained by the scarcity of missense variants with only 36-38 nonsynonymous mutations detected in 11 subunits at the continental scale and mutation rates as low as 0-0.0069 mutation per nucleotide. In our continentwide survey, the α5 subunit emerged as the most conserved nAChR gene, with no detectable nonsynonymous mutations in *An. gambiae* and *An. coluzzii* populations. The pronounced conservation of this subunit is supported by previous studies (Perry et al. 2021; Lu et al. 2022). To our knowledge, no amino acid substitutions have been reported in α5 in insect pests, even among species with well-documented insecticide resistance profiles (Matsuda et al. 2020). Comparatively, other highly conserved Cys-loop receptors, such as the GABA receptor gene *rdl*, do harbor nonsynonymous mutations in *Anopheles*, including substitutions that arose and spread within the last 70 years in response to dieldrin pressure (ffrench-Constant et al. 1993; Thompson et al. 1993; Kamdem et al. 2017; Grau-bové et al. 2020). Standing variation identified across nAChRs in this study differs from resistance mutations often detected in the gene family among agricultural pests in many ways. First, most resistance-associated mutations concentrate on the β1 subunit and to a lesser extent α6 and α3 whereas standing variation was detected in all subunits except α5 in *An. gambiae* and *An. coluzzii* (Liu et al. 2005; Bass et al. 2011; Puinean et al. 2013; Chen et al. 2017; Yin et al. 2024; Cheng et al. 2025). Amino acid changes were also more frequent in α3, α9, and β2 (with six, seven, and nine nonsynonymous mutations respectively) which are rarely implicated in resistance in agricultural pests.

Our data which were collected from wild *An. gambiae* and *An. coluzzii* populations in the absence of neonicotinoid or spinosyn pressure indicate the effects of purifying selection acting on constrained loci leaving only rare nonsynonymous variants with limited potential to alter receptor function. Precisely, resistance-associated substitutions typically occur in orthosteric or allosteric binding sites of nAChR subunits, exert strong phenotypic effects, and have fitness implications in pest species (Erdmanis et al. 2012; Puinean et al. 2013; Perry et al. 2021; Cheng et al. 2025). In contrast, most amino acid substitutions in our study lie outside of the protein’s functional domains and tend to conserve the biochemical properties of the original residues. Notably, no missense mutations were found in any of the four transmembrane helices (TM1–TM4), and only one low-frequency substitution (T220P in α3) fell within a primary ligand-binding loop (loop C). The D202E in α6 occurred in loop F, but loop D-F in α subunits play only a secondary role in stabilizing the ligand-binding pocket (Jones and Sattelle 2010). This spatial distribution is consistent with well-known constraints in Cys-loop receptors, where loops form the orthosteric binding pocket and the transmembrane helices mediate channel gating and desensitization. Amino acid residues in these domains are subjected to enhanced constraints because they contribute to the extracellular–transmembrane interface where general properties such as shape and relative positioning of the opposing surfaces determine Cys-loop receptor function (Tsunoyama and Gojobori 1998; Cymes and Grosman 2021; Godellas et al. 2024).

SIFT scores, which measure the deleteriousness of amino acid replacements also supported that standing variation in constrained nAChRs is represented by mutations predicted to be tolerated (Ng and Henikoff 2001). Most of these variants especially high-frequency nonsynonymous mutations detected in *An. gambiae* and *An. coluzzii* have SIFT scores > 0.05 indicating tolerated substitutions. Conservative substitutions are unlikely to disrupt protein structure or receptor–ligand interactions and may thus persist in *Anopheles* populations due to their minimal fitness costs (Somers, Luong, Mitchell, et al. 2017; Somers, Luong, Batterham, et al. 2017; Zhang et al. 2022).

### 4.2 Low-frequency missense variants reflect mutation–selection balance in nAChRs

A second line of evidence supporting the hypothesis that *An. gambiae* and *An. coluzzii* populations tolerate only low-impact mutations in nAChR subunits comes from the frequency distribution of missense variants. In contrast to resistance mutations that can be rapidly driven to high frequencies by positive selection (Perry et al. 2025: 25; Sparks et al. 2025), most missense variants identified in our dataset were rare and likely recessive changes masked in heterozygous individuals (Dussex et al. 2023). Specifically, mutant homozygotes were observed in less than 5% of individuals for most variants, and 22 out of the 36 missense substitutions were never found in the homozygous state. This skew toward heterozygote-only variants is consistent with a mutation–selection balance model (Haldane 1937), in which mildly deleterious alleles are maintained at low frequency in heterozygotes but are purged from the population when homozygous due to their detrimental fitness effects (Schmenger et al. 2022; Dussex et al. 2023; Smeds and Ellegren 2023).

One important exception is the α6 D202E substitution, which has risen to high frequency in *An. gambiae*. However, haplotype-based tests provide no evidence for a recent selective sweep surrounding this variant. Extended haplotype homozygosity (EHH) decayed rapidly from ∼1.0 to < 0.1 within just 200 bp on either side of the D202E site and was indistinguishable between wildtype and mutant sequences across a 10-kb window. The absence of extended haplotype structure suggests that the rise in frequency of D202E was not driven by recent directional selection, which would be expected to produce long haplotypes of reduced diversity through hitchhiking with surrounding loci (Maynard-Smith and Haigh 1974; Sabeti et al. 2002). In addition, at a finer geographic scale within Cameroon, we found that genotype frequencies showed no variation across eco-geographic regions suggesting that D202E is not subject to environmentally driven selection. However, the frequency of the derived allele was lower in the Kisumu *An. gambiae* colony than in conspecific Cameroonian field populations, suggesting that selection, founder effects, colony drift, or bottlenecks shape a distinct evolutionary trajectory for α6 mutations in laboratory conditions. These findings underscore the importance of tracking haplotype variation in both field and laboratory populations to better understand how evolutionary and ecological forces modulate nAChR diversity and insecticide target-site evolution.

### 4.3 Emergence of α6 haplotypes within the *An. gambiae* complex

Despite strong purifying selection across nAChRs, the a6 subunit stands out as an exception – harboring two high-frequency substitutions (I198M and D202E) with the derived alleles fixed in *An. gambiae* but absent in *An. coluzzii*. Both variants define species-specific haplotypes with variable spinosad susceptibility. While other nAChR subunits such as α7 and β1 also exhibit species-level clustering based on synonymous variation, only α6 shows nonsynonymous divergence, highlighting exon 7 as a key region for functional variation. α6 is unique among nAChRs in that loss-of-function mutants in many insect species are viable and exhibit spinosyn resistance without severe developmental defects (Perry et al. 2007; Perry et al. 2021; Lu et al. 2022; Martelli et al. 2022). This reduced fitness cost may allow *An. gambiae* α6 to evolve under more relaxed constraints, accumulating conservative substitutions that subtly modulate receptor function (Perry et al. 2025; Sparks et al. 2025). The presence of the derived M198–E202 haplotype in both *An. gambiae* and *An. arabiensis*, but not in *An. coluzzii*, suggests possible introgression between *An. gambiae* and *An. arabiensis* or ancient polymorphism followed by lineage sorting (Fontaine et al. 2015). Frequency distributions in more distant *Anopheles* lineages support the idea that at least two derived haplotypes have recently spread throughout the *An. gambiae* complex. Other insecticide targets such as the voltage-gated sodium channel (*Vgsc*), the dieldrin resistance gene (*Rdl*), and the acetylcholinesterase (*AChE*) locus have shown a similar evolutionary trajectory within this species complex. In these genes, selection from standing genetic variation first contributed to the emergence of resistance alleles in one taxon which were subsequently spread to sibling species through introgression as insecticide pressure increased (Weill et al. 2000; Djogbénou et al. 2008; Clarkson et al. 2014; Norris et al. 2015; Assogba et al. 2016; Grau-bové et al. 2020; Clarkson et al. 2021). Consequently, while the emerging α6 haplotypes show no evidence of recent positive selection, their functional effects and frequencies should be evaluated and monitored during insecticide rollout.

Our study highlight variability in baseline susceptibility to Spinosad – a spinosyn insecticide targeting the nAChR α6 subunit – among α6 haplotypes (Perry et al. 2007; Perry et al. 2021; Perry et al. 2025). *An. coluzzii* larvae, fixed for the ancestral I198–D202 haplotype, exhibited rapid and complete mortality at field-realistic doses of spinosad, with minimal pupation. In contrast, *An. gambiae* (Nkolondom population), fixed for the derived M198–E202 haplotype, displayed delayed mortality and substantially higher pupation rates (57% at 0.1 ppm; 3% at 0.5 ppm), suggesting enhanced tolerance. A broader sampling of *An. gambiae* and *An. arabiensis* as well as experimental validation are still needed to clearly establish a link between the derived M198–E202 haplotype and susceptibility to spinosad. Moreover, the mechanisms underpinning this apparent phenotypic advantage remains unclear. Both I198M and D202E are SIFT-tolerated and predicted to have minimal functional impact. Importantly, both mutations lies around loop F, ∼40 Å from the proposed spinosad allosteric site (Puinean et al. 2013). However, static distance estimates do not capture dynamic determinants of spinosyn action such as gating kinetics, desensitization, inter-subunit coupling, and long-range allostery (Thompson et al. 2010; Gulsevin 2020). Thus, these substitutions could modulate conformational equilibria or coupling without directly affecting ligand-receptor interactions.

It is worth noting that the most widespread spinosad resistance mutation in agricultural pests is a glycine-to-glutamic acid substitution (G275E) in α6 (Puinean et al. 2013; Perry et al. 2025; Sparks et al. 2025). Although this mutation is not structurally analogous to the D202E substitution found in *An. gambiae*, the presence of a derived glutamic acid residue in both mutations raises the possibility that glutamic acid substitutions at key positions within α6 could alter spinosyn susceptibility. CRISPR-mediated knockouts and site-directed mutagenesis targeting I198M and D202E in *Anopheles* offer promising tests to unravel the physiological and toxicological consequences of these substitutions (Watson et al. 2010; Puinean et al. 2013; Perry et al. 2021). Heterologous expression using *Xenopus* oocytes has been applied to test the affinity of *An. gambiae* nAChR subtypes containing α1, α2, α3, α8 and β1 subunits to neonicotinoids (Ito et al. 2024). Extending these approaches to α6-containing subtypes would provide insights into the role M198–E202 haplotype in modulating spinosad sensitivity.

### 4.4 Implications for vector control and resistance management

Resistance management strategies such as insecticide rotations rely on the assumption that the frequency of resistance alleles will gradually decline once the selective agent is removed from the environment (WHO 2012). However, empirical evidence increasingly challenges this premise and shows that resistance alleles can sometimes be stably maintained in wild populations even in the absence of active selection. For instance, *rdl* mutations persist in *Anopheles* populations across Africa, and the frequency of *kdr* mutations in the Vgsc gene remains high in *Aedes aegypti* from Brazil, decades after the cessation of associated insecticide use (Kamdem et al. 2017; Miles et al. 2017; Macoris et al. 2018; Grau-bové et al. 2020; Boddé et al. 2025). As a result, surveillance of standing genetic variation at target sites could inform proactive management of resistance alleles for new classes of insecticides introduced into vector control. Competitive or allosteric modulators of nAChRs including neonicotinoids, butenolides and spinosyns are progressively being introduced in malaria vector control interventions (Hertlein et al. 2010; WHO 2010; WHO 2025). In addition to public health interventions, widespread agricultural use of neonicotinoids has often raised concerns about the role of residual exposure and cross-resistance in driving the emergence of resistance in *Anopheles* (Georghiou 1972; Lines 1988; Chouaibou et al. 2019; Ambadiang et al. 2024; Tchigossou et al. 2024). The absence of selective sweeps at nAChR loci, and the low frequency of heterozygote-exclusive nonsynonymous variants suggests that target-site resistance mutations have not yet been selected by pesticide pressure in the environment. However, a recent report of clothianidin resistance in *An. funestus*—emerging within just two years of neonicotinoid rollout in Uganda—reinforces the need to monitor molecular changes at target sites (Kamya et al. 2024; Oruni et al. 2025). Thus, the comprehensive survey of genetic variation across 11 nAChR subunits in *An. gambiae* sensu lato populations provides baseline data for tracking the emergence of resistance alleles as these compounds are deployed. Our findings also lay the groundwork for characterizing nAChR variation across all major *Anopheles* vectors, many of which remain poorly studied in the context of vector control.

## Acknowledgements

This study was supported by the National Institutes of Health (R01AI150529 to C.K.). We thank our field assistants in Cameroon for their help with larval sampling and collections.

## Data availability statement

Variant call format (VCF) files from the *Anopheles gambiae* 1000 Genomes Project analyzed in this study are available from the MalariaGEN repository (https://www.malariagen.net/project/ag1000g/). VCFs for *Anopheles arabiensis*, *An. melas*, *An. quadriannulatus*, and *An. merus* are available from the European Nucleotide Archive (ENA) under BioProject PRJNA1331534. Accession numbers are provided in Supplementary Tables S1 and S2.

## Author contributions

Designed research: CF, CK; performed research: CF, DR, FA, MA, MP, CH, CK; analyzed data: CF, CK; wrote the paper: CF, CK.

## Supplementary material

**Table S1. City and country of origin for 532 *Anopheles gambiae* and *An. coluzzii* specimens from the *Anopheles gambiae* 1000 Genomes Project**

**Table S2. Genotypes at α6 I198M and D202E for four species in the *Anopheles gambiae* complex**

